# Development of a CRISPR-Cas9 genome editing system for the extremely thermophilic bacterium *Anaerocellum bescii*

**DOI:** 10.64898/2026.10.26.761776

**Authors:** Joey L. Galindo, Jonathan M. Conway

**Author notes:** **Correspondence:** Jonathan M. Conway.

## Abstract

*Anaerocellum* (f. *Caldicellulosiruptor*) *bescii* is an extremely thermophilic (T_opt_ 75-78 ℃) anaerobic bacterium and is also a highly efficient degrader of lignocellulose, making it a promising platform for metabolic engineering. To date, chromosomal genetic manipulations in *A. bescii* have required multistep procedures involving homologous recombination with integrating vectors and counterselection. Here, we demonstrate the first implementation of CRISPR-Cas9 genome editing in *A. bescii* facilitated by a thermostable Cas9 enzyme from *Geobacillus stearothermophilus* (GeoCas9). Delivery of heterologous CRISPR machinery via a replicating vector enabled us to individually and sequentially delete genes for lactate dehydrogenase (*ldh*) and *pyrE* in *A. bescii*. We then deployed this system to delete a novel target, tryptophan synthase (*trpBA*), generating a tryptophan auxotrophic strain of *A. bescii*. Using this strain, we show *trpBA* complementation enables selection and maintenance of a replicating vector expressing a heterologous enzymatic reporter gene.

**Importance:** Extreme thermophiles, especially species that can naturally break down recalcitrant lignocellulose, such as *Anaerocellum* (f. *Caldicellulosiruptor*) *bescii*, represent compelling chassis for next-generation bioprocessing, offering the capability to convert inexpensive and plentiful renewable feedstocks into fuels and industrial chemicals. However, engineering these non-model organisms for bioprocessing applications requires robust genetic engineering tools, as techniques developed in mesophilic model organisms often translate poorly. CRISPR-Cas genome editing, now widely used in biotechnology, is one such tool that faces limitations at the extreme native temperatures of these organisms. Here we establish a CRISPR-Cas9-based genome editing system in *A. bescii*, expanding the genetic toolbox available for this extremely thermophilic bacterium. By streamlining targeted genetic modifications, this approach has the potential to accelerate the construction of engineered strains for metabolic engineering and bioprocess applications.

## Introduction

*Anaerocellum* (formerly *Caldicellulosiruptor*) *bescii* is the most thermophilic lignocellulose-degrading bacterium known, with an optimal growth temperature of 75-78 ℃ (1–3). Its strong lignocellulose-degrading capacity is enabled by a large repertoire of multi-domain enzymes that efficiently deconstruct a wide variety of recalcitrant lignocellulosic biomass (3–5). These unique characteristics make *A. bescii* an attractive chassis for synthetic biology and novel bioprocessing approaches (2, 4, 6). As such, *A. bescii* has been engineered to produce several non-native products of commercial relevance, including ethanol, acetone, and 2,3-butanediol (7–11). These metabolic engineering efforts were enabled by the development of a functional genetic system in *A. bescii*, most notably the implementation of a highly thermostable kanamycin resistance (*htk*) marker for selection on kanamycin, and generation of uracil auxotroph strains (Δ*pyrE* or Δ*pyrF*) for positive selection via complementation and counterselection on 5-FOA (12–15). Subsequent deletion of the lactate dehydrogenase (*ldh*) gene further enhanced engineering work in *A. bescii* by eliminating lactate formation and redirecting metabolic flux toward formation of desirable products (12, 15). Yet targeted chromosomal genetic manipulations in *A. bescii* have relied on multistep homologous recombination procedures, which, although effective, is often laborious and time-consuming, requiring transformation with an integrative vector (selected by kanamycin resistance or uracil prototrophy) followed by 5-FOA counterselection to resolve the initial merodiploid intermediate (2, 12, 15). Furthermore, strains derived from an early 5-FOA-selected uracil auxotroph background strain exhibited extensive mutations and transposon insertions associated with genome instability and unwanted phenotypic changes (16).

In contrast, CRISPR-Cas based genome editing offers a more straightforward approach for genetic manipulation. CRISPR-Cas systems fall into two broad classes, which have been further subdivided into seven major types (17–19). Class 1 systems utilize multiple effector Cas proteins, while class 2 systems consist of a single effector Cas protein, making this latter category ideal for exogenous delivery and expression in non-native hosts (18–20). Class 2 CRISPR systems include the Type II-Cas9 systems that have become ubiquitous in biotechnology, archetypically represented by the Cas9 from the bacterium *Streptococcus pyogenes* (SpCas9) (18, 19, 21). In natural CRISPR-Cas9 systems, a target-specific CRISPR RNA (crRNA) first hybridizes to a short transactivating RNA (tracrRNA), after which the resulting duplex binds to the Cas9 and directs it to cleave the complementary target DNA sequence (19, 21). However, as a tool for genetic editing, the crRNA and tracrRNA are typically fused into a single guide RNA (sgRNA) (21, 22). Delivery or expression of this guide RNA (gRNA) and the Cas9 effector within the target host produces a dsDNA break at the target so long as the target sequence is adjacent to a specific protospacer adjacent motif (PAM) sequence (19, 20, 22). Repair of this dsDNA break can occur through error-prone non-homologous end joining (NHEJ) or homology-directed repair (HDR) using a supplied DNA repair template (19–21), although many bacteria lack the machinery for NHEJ (23, 24).

Cas effector proteins derived from mesophiles, such as SpCas9, are not nearly thermostable enough to function effectively in a thermophilic bacterium like *A. bescii* (20, 25). Fortunately, a number of thermostable Cas9 variants have been characterized, such as those from *Geobacillus stearothermophilus* (GeoCas9), *Geobacillus thermodenitrificans* (ThermoCas9), *Geobacillus sp.* LC300 (CaldoCas9), *Ignavibacterium* (IgnaviCas9), *Anoxybacillus flavithermus* (anoCas9), *Acidothermus cellulolyticus* (AceCas9) and *Alicyclobacillus tengchongensis* (AtCas9) (26–32). In turn, several of these Cas9s have been successfully deployed as tools for genome editing in non-native thermophilic bacterial hosts, including in *Acetivibrio thermocellus* (f. *Clostridium thermocellum*), *Thermoanaerobacter kivui*, *Thermoanaerobacter ethanolicus*, *Bacillus smithii*, and *Thermus thermophilus* (27, 30, 33–36). GeoCas9 in particular has been used successfully for CRISPR editing in several of these species (33, 34, 36). *Harrington et al.* previously determined GeoCas9 utilizes a PAM sequence of 5’-NNNNCRAA-3’ and has an optimal spacer length of 22 nt (26). To create a single guide RNA, they joined this spacer to a repeat sequence fused to its associated tracrRNA via a GAAA tetraloop, allowing for *in vitro* nuclease activity at temperatures up to 70 ℃ (26). Given this thermostability and its track record in other thermophilic bacteria, we reasoned that GeoCas9 could be adapted for genome editing in *A. bescii*.

In this study, we develop a pipeline for rapid genetic knockouts in *A. bescii* mediated by GeoCas9. Delivery of sgRNAs, the *GeoCas9* gene, and repair templates via a replicating vector enabled individual knockouts of the previously targeted *ldh* and *pyrE* genes in *A. bescii* without the need for counterselection on 5-FOA. We achieved sequential editing by curing the Δ*pyrE* strain of its CRISPR knockout plasmid and transforming it with a second knockout plasmid targeting *ldh*. We then demonstrate the versatility of this system by knocking out a novel target, tryptophan synthase (*trpBA*), generating a tryptophan-auxotrophic strain of *A. bescii*. We further show that restoration of tryptophan prototrophy through *trpBA* complementation enables selection of *A. bescii* transformants carrying a replicating vector expressing the *trpBA* cassette and also containing a heterologous enzymatic reporter gene. Taken together, this work describes an accelerated workflow for genetic knockouts in *A. bescii,* representing the first implementation of CRISPR-Cas9 editing in this extremely thermophilic bacterium.

## Results and Discussion

### Strategy for CRISPR-Cas9 editing in *A. bescii*

To deliver the components needed for CRISPR-Cas9 editing in *A. bescii* our approach relied upon an established replicating vector backbone that contains the native *A. bescii* plasmid (*pAthe02*) and the *htk* gene, which respectively enable replication in *A. bescii* and selection on kanamycin (12, 13). CRISPR machinery, including the codon optimized *GeoCas9* gene, sgRNA expression constructs, and repair-templates were added onto this backbone (**Fig. 1**). We expressed the GeoCas9 with the relatively low strength *Athe_1331* promoter (P_1331_), surmising that this lower level of expression (37), combined with borderline thermostability of GeoCas9 (26) at the typical 70-75 ℃ culture temperatures of *A. bescii*, would limit any potential toxicity. Initially, we tested the viability of expressing GeoCas9 alone by transforming *A. bescii* with a replicating vector (pJLG236) containing the P_1331_-*GeoCas9* cassette but no gRNA (**Fig. 1**). After successful transformation and confirmation of the correct *GeoCas9* sequence (**Fig. S1a**), this strain was designated JGAB51 and would serve as the no gRNA control strain in subsequent implementation of knockout vectors with gRNAs. A full list describing the strains constructed in this study, including JGAB51, can be found in **Table 1**.

**Figure 1.**
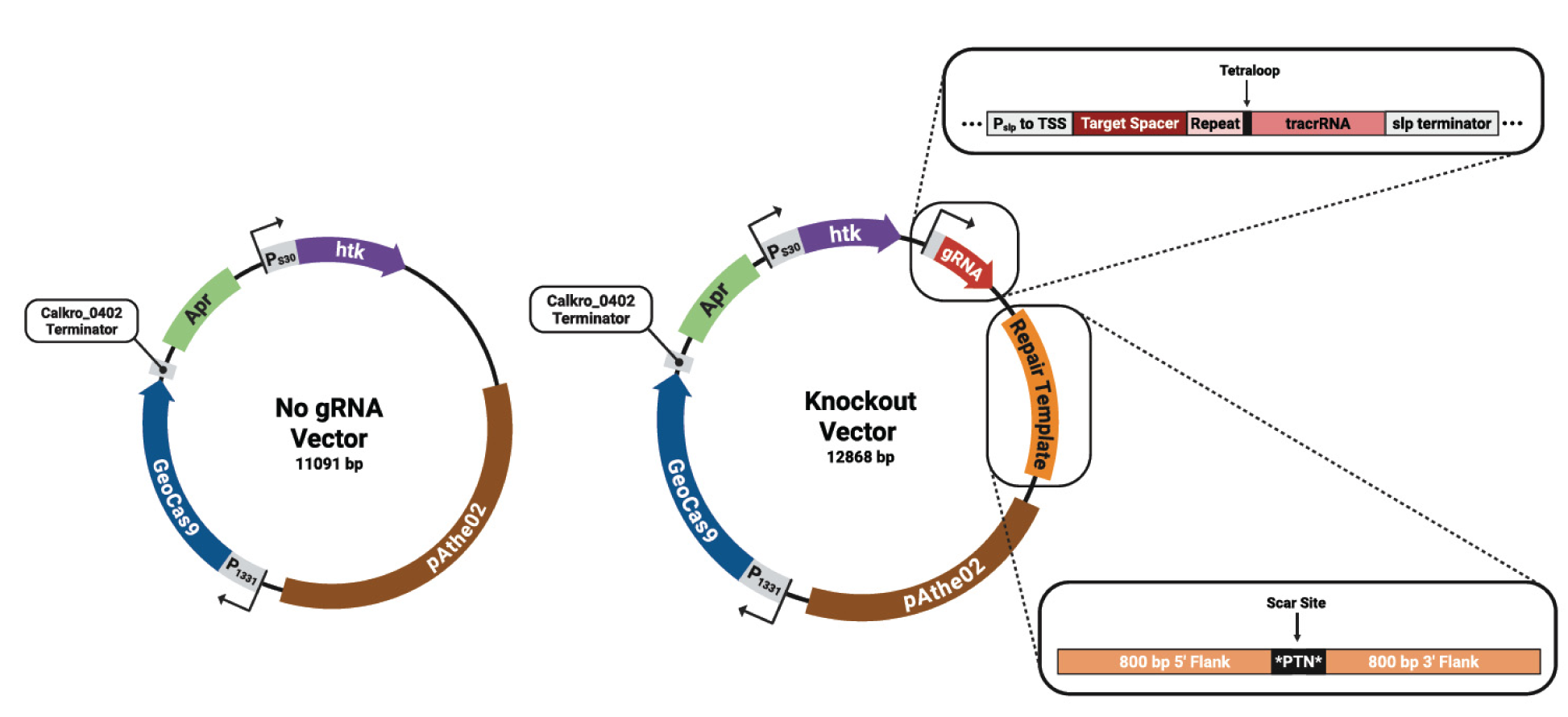
Plasmid maps of GeoCas9-expressing replicating vectors transformed into *A. bescii*. These vectors contain the gene for a thermostable Cas9 from *Geobacillus stearothermophilus* (GeoCas9) expressed by the *Athe_1331* promoter (P_1331_). The left no gRNA vector (pJLG236) served as the control plasmid lacking a sgRNA, while knockout vectors (pJLG240-41 & pJLG250) on the right contain gene specific sgRNA sequences expressed by the s-layer protein promoter (P_slp_) truncated to its transcriptional start site (TSS). To facilitate homology directed repair (HDR), these knockout vectors also contain repair templates with a unique *PTN* (tagccaacgaattaa) scar site. Created in BioRender. https://BioRender.com/jy32y1f.

**Table 1.** *A. bescii* strains constructed and used in this study.

| Strain Designation | Parent | Transforming Plasmid | Description |
| --- | --- | --- | --- |
| DSM 6725 (wild type) | n/a | n/a | n/a |
| JGAB51 | DSM 6725 | pJLG236 | no gRNA control |
| JGAB53 | DSM 6725 | pJLG240 | $\Delta ldh$ (*Athe_1918*) |
| JGAB54 | DSM 6725 | pJLG241 | $\Delta pyrE$ (*Athe_1382*) |
| JGAB55 | JGAB54 | pJLG240 | $\Delta pyrE \Delta ldh$ |
| JGAB56 | DSM 6725 | pJLG250 | $\Delta trpBA$ (*Athe_1690-91*) |
| JGAB57 | JGAB56 | pJLG253 | $\Delta trpBA$ pJLG253[*trpBA*, *Cmβgal*] |

Having established that *A. bescii* could maintain the *GeoCas9* gene on a replicating vector without a gRNA, we next tested knockout vectors containing gRNAs. These gRNAs were expressed from a modified version of the strong, constitutive s-layer protein promoter (P_slp_). For gRNA expression, this typically 200 bp promoter was truncated at its 3’ end to the previously determined transcriptional start site, 74bp upstream of the native start codon (37), removing the ribosome binding site and enabling transcription of gRNAs without additional 5’ nucleotides. Based on the functional characterization of GeoCas9 by *Harrington et al.* (26), we designed our sgRNAs with a 22 nt spacer and the same repeat-tetraloop-tracrRNA sequence (**Fig. 1** & **Table S1**). Because *A. bescii* lacks identifiable machinery for NHEJ (23, 24, 38), each knockout plasmid also contained a repair template to facilitate HDR, consisting of two 800bp homology arms flanking the targeted genomic region (**Fig. 1** & **Supplementary Table S1**). A 15 bp scar sequence, *PTN* (tagccaacgaattaa), was inserted between the homology arms to track repair-template incorporation into the genome (**Fig. 1**). Sequences for various CRISPR components, including the repeat-tetraloop-crRNA, spacer sequences and associated PAM sites, repair templates, and the codon optimized *GeoCas9* gene, are provided in **Supplementary Table S1**.

### Implementation of GeoCas9 mediated genetic knockouts and strain construction

Two target genes were initially chosen to test the CRISPR-Cas9 knockout vector design: *ldh* (*Athe_1918*) and *pyrE* (*Athe_1382*), which respectively encode lactate dehydrogenase and orotate phosphoribosyl transferase enzymes (38). As discussed above, both genes have previously been deleted through homologous recombination in *A. bescii* due to the utility of the resulting strains for further genetic manipulations and metabolic engineering (12, 15). Importantly, these genetic deletions result in clear phenotypes: loss of *ldh* eliminates lactate production, while deletion of *pyrE* causes uracil auxotrophy (12, 15). Using these targets, we developed a post-transformation workflow for CRISPR-Cas9-facilitated strain construction in *A. bescii* (**Fig. 2a**).

**Figure 2.**
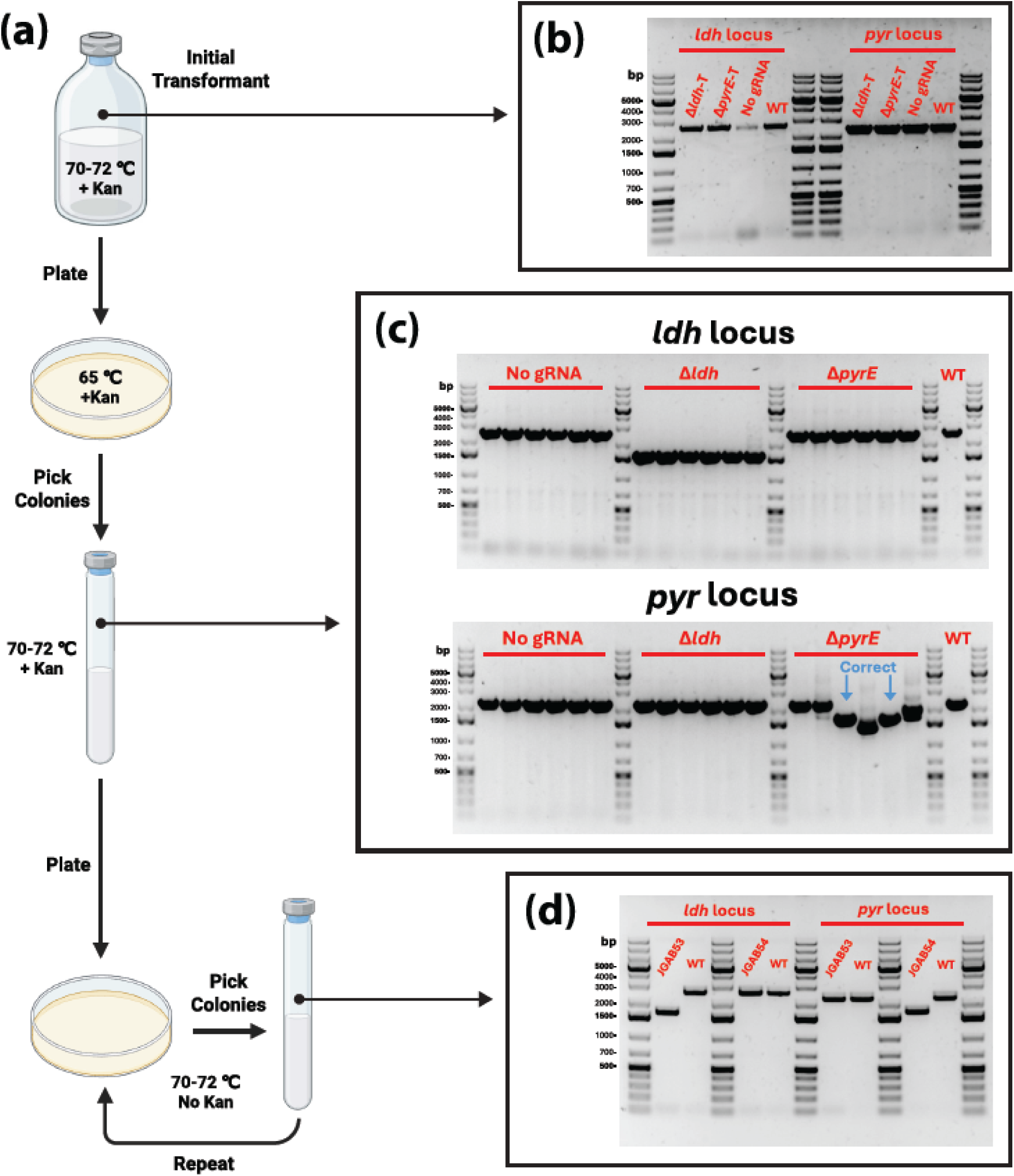
CRISPR-Cas9 mediated genetic knockouts of *ldh* and *pyrE* in *A. bescii*. **(a)** Diagram of developed strain construction workflow using GeoCas9. **(b)** Colony PCR of initial successful transformant cultures (Δ*ldh*-T & Δ*pyrE*-T) prior to plating, as well as the no gRNA control strain (JGAB51), and wild type *A. bescii* (WT) gDNA. **(c)** PCR of *A. bescii* colonies picked after initial plating on selective media as well as WT gDNA. Blue arrows indicate which sequenced Δ*pyrE* colonies matched the repair template. **(d)** Colony PCR of final sequence confirmed *A. bescii* strains containing the *ldh* and *pyrE* deletions, designated JGAB53 and JGAB54 respectively. For panels **b-d**, expected amplicons from PCR on the locus surrounding *ldh* (**Table S2**; Primers JLG288 & 289) were 2.6 kb (WT) and 1.7 kb (Δ*ldh*), while those targeting *pyrE* (**Table S2**; Primers JLG290 & 291) were 2.3 kb (WT) and 1.7 kb (Δ*pyrE*). Panel **a** was created in BioRender. https://BioRender.com/r33j2as.

For the initial transformation of the Δ*ldh* (pJLG240) and Δ*pyrE* (pJLG241) CRISPR vectors, electroporation of competent cells was followed by recovery and subsequent selection with kanamycin in liquid media at 70-72 ℃. PCR analysis from the initial transformant cultures yielded amplicons corresponding to the wild-type *pyr* and *ldh* loci, indicating genome editing had not yet occurred (**Fig. 2b**). This is consistent with the previous findings of *Harrington et al.* who showed that while GeoCas9 can function *in vitro* at up to 70 ℃, its activity drastically decreases above 65 ℃ (26). Thus, to promote GeoCas9 activity, transformant cells were grown at a lower temperature of 65 ℃ when first plated in solid selective media (**Fig. 2a**). Colonies that were picked and screened via PCR after this plating step had deletions to the desired target gene (*ldh* or *pyrE*). Colonies of JGAB51 (no gRNA) that were plated alongside the strains transformed with gRNA-containing vectors showed no cutting at either locus (**Fig. 2c**). Of the 6 colonies initially picked for cells transformed with the Δ*ldh* CRISPR plasmid (pJLG240), all had the desired genomic deletion (**Fig. 2c**). For the Δ*pyrE* CRISPR plasmid (pJLG241) cells, the results were mixed, with only 2 out of the 6 having the correct amplicon, while other colonies showed either no cutting or deletions of Δ*pyrE* that were different from the provided repair template (**Fig. 2c**). Subsequent sequencing of these PCR products confirmed that colonies with the expected amplicon size for their respective knockout target had genotypes consistent with HDR using the provided template, containing deletions that matched the flanking regions as well as incorporation of the *PTN* scar site in the location of the target gene.

To complete construction of the edited *A. bescii* strains, cells were cured of their CRISPR plasmids by plate-purifying colonies with the desired deletions two additional times on non-selective media lacking kanamycin, after which the genomic deletions were confirmed again by sequencing (**Fig. 2a & d**). Curing cells of their CRISPR knockout plasmids through cycles of outgrowth on non-selective media allows these strains to be transformed again with other plasmids that use the kanamycin resistance marker, while also removing the *GeoCas9* gene. As in previous *A. bescii* studies that used homologous recombination, plate purification also helps ensure a homogeneous population with the desired deletion. Plate purified strains were then tested for growth in liquid kanamycin selective media, none of which grew after 40 hours of incubation at 70 ℃. The resulting Δ*ldh* and Δ*pyrE* strains were designated JGAB53 and JGAB54 respectively (**Table 1**).

Next, we experimentally confirmed that these strains displayed the expected phenotypes: abolition of lactate production in JGAB53 (Δ*ldh*), and uracil auxotrophy in JGAB54 (Δ*pyrE*). To test the former, JGAB53 and wild type *A. bescii* were grown in triplicate in defined cellobiose medium (DC516) for 30 hours at 70 ℃, after which the major natural fermentation products, acetate and lactate, were quantified by HPLC (**Fig. 3a**). As expected, no lactate was detected in cultures of JGAB53, while wild type *A. bescii* produced 8.7 mM lactate. JGAB53 also produced increased acetate titers of 4.6 mM compared to 1.8 mM in cultures of wild type cells. This increase in acetate production is consistent with previous observations of other Δ*ldh A. bescii* strains, as the loss of *ldh* eliminates a major metabolic outlet and requires flux be redirected elsewhere (15). Uniquely, JGAB53 is the first strain of *A. bescii* carrying only the Δ*ldh* deletion, whereas previous Δ*ldh* strains required a *ΔpyrE or ΔpyrF* background to enable 5-FOA counterselection during homologous recombination-based editing (12, 15). We then verified that JGAB54 was an uracil auxotroph by growing it in triplicate, alongside wild-type cells, on DC516 medium with or without the addition of 20 µM uracil. While wild-type *A. bescii* was able to grow on both types of defined media, JGAB54 grew only in the presence of uracil, with no growth detected on medium lacking uracil after 72 hours at 70°C (**Fig. 3b**).

**Figure 3.**
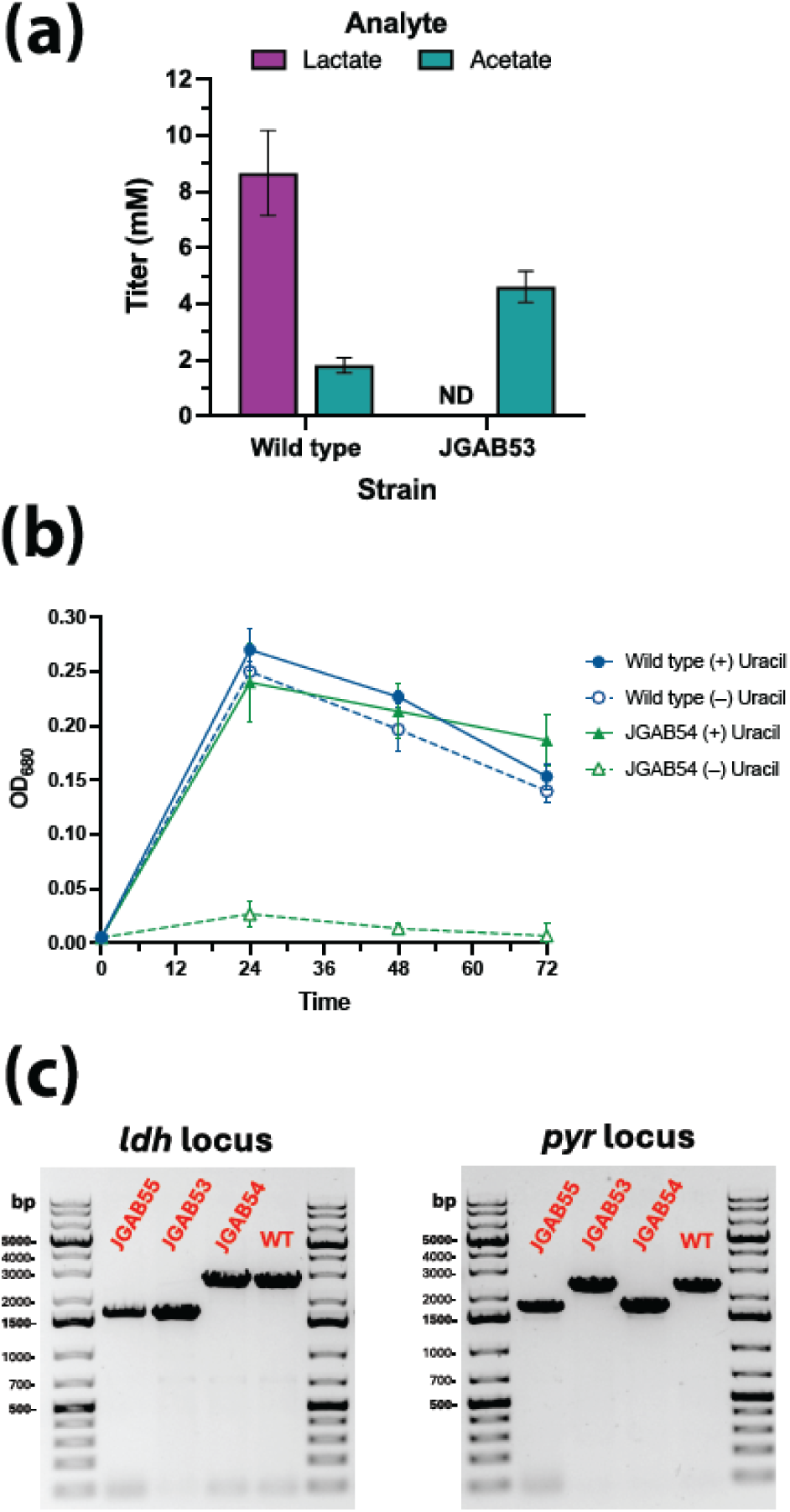
Characterization of *A. bescii* knockout strains constructed via CRISPR-Cas9 genome editing. **(a)** Fermentation products (acetate and lactate) quantified via HPLC of wild type and JGAB53 (*Δldh*) *A. bescii* cultures grown for 30 hours on defined cellobiose media (DC516) at 70 ℃. ND=not detected. **(b)** 72-hour growth curves of wild type and JGAB54 (*ΔpyrE*) *A. bescii* grown on DC516 media with or without 20 µM uracil at 70 ℃. Error bars in panels **a** & **b** represent one standard deviation from triplicate biological replicates. **(c)** Final colony PCR of JGAB55 (*ΔpyrE Δldh*), constructed by transforming JGAB54 with the *Δldh* CRISPR knockout vector (pJLG240), alongside gDNA from WT *A. bescii*, JGAB53, and JGAB54. Expected amplicons for PCR on the *ldh* locus (**Table S2**; Primers JLG288 & 289) were 2.6 kb (WT) and 1.7 kb (Δ*ldh*), while those for the *pyrE* locus (**Table S2**; Primers JLG290 & 291) were 2.3 kb (WT) and 1.7 kb (Δ*pyrE*).

Having established the phenotypes for both individual deletions, we next applied this CRISPR editing method to generate a strain with multiple sequential knockouts through transformation with a different plasmid. Specifically, JGAB54 (Δ*pyrE*) was transformed with the Δ*ldh* plasmid (pJLG240) to generate a Δ*pyrE* Δ*ldh* strain. Successful transformants underwent the same process as described above, with 5 of the 6 colonies initially picked having the desired *ldh* deletion (**Fig. S1b & c**). Following the plasmid curing process, sequences matching the repair template at both the *pyrE* and *ldh* loci were confirmed, each containing the *PTN* scar site; the resulting Δ*pyrE* Δ*ldh* strain was designated JGAB55 (**Table 1**). Together, these results demonstrate that the GeoCas9-based CRISPR system supports sequential gene deletions in *A. bescii* through repeated rounds of plasmid transformation, plating at 65°C, and plasmid curing.

### Leveraging CRISPR-Cas9 genome editing to establish tryptophan-based selection

Next, we aimed to apply this GeoCas9-based system to the tryptophan biosynthesis locus in *A. bescii*, with the goal of generating an auxotrophic strain that could enable a new tryptophan-based selection strategy in this organism. Tryptophan is a large and energetically costly amino acid to synthesize, and its biosynthetic pathway is highly conserved across multiple domains of life (39–41). Tryptophan biosynthesis comprises five enzymatic steps that convert chorismate, generated via the shikimate pathway, into L-tryptophan (40, 41). In bacteria and archaea, the genes encoding this pathway are often clustered within a 5-7 gene operon (39, 40). The *A. bescii trp* locus comprises seven genes (*Athe_1690–1696*), organized as *trpEGDCFBA*, in which *trpEGD* and *trpCFBA* form two blocks of overlapping open reading frames (**Fig. 4a**) (38). Tryptophan prototrophy has previously been used for selection in other thermophilic microbes, including the archaeon *Thermococcus kodakarensis* and the bacterium *Thermus thermophilus* (2, 42, 43). We therefore targeted the *trpBA* genes at the end of the *A. bescii* tryptophan biosynthesis locus for deletion, aiming to generate a tryptophan-auxotrophic strain and subsequently establish a new selection strategy through *trpBA* complementation.

**Figure 4.**
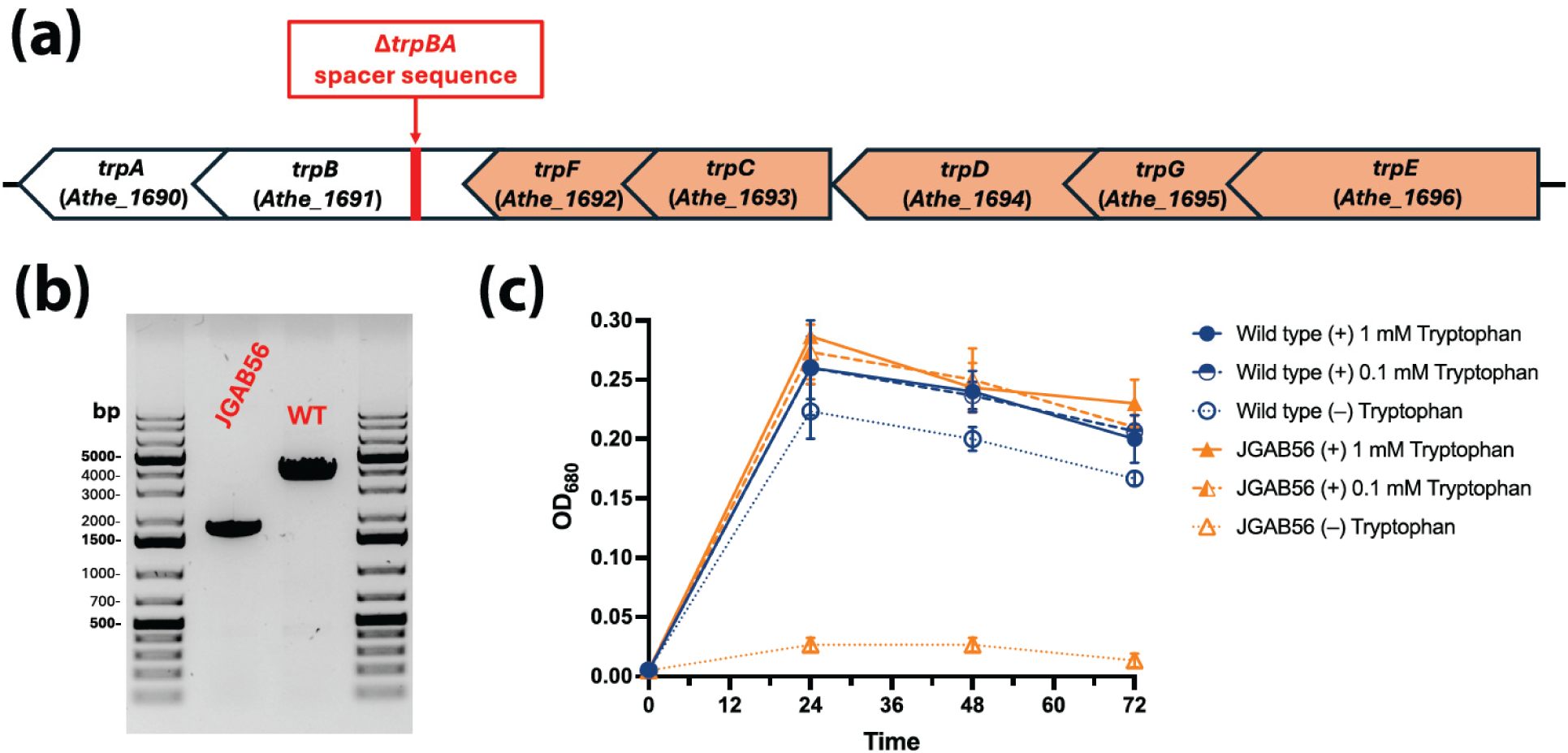
Knockout of the tryptophan synthase genes (*trpBA*; *Athe_1690-91*) in *A. bescii*. **(a)** Diagram of the 7 gene tryptophan (*trp*) synthesis locus in the *A. bescii* genome. *trpBA* (colored in white) was targeted for CRISPR-CAS9 mediated deletion. **(b)** PCR of final plate purified, sequence confirmed *ΔtrpBA* strain, designated JGAB56 alongside wild type (WT) *A. bescii* gDNA. Expected amplicons sizes around the *trpBA* locus (**Table S2**; Primers JLG319 & 320) were 3.7 kb (WT) and 1.7 kb (Δ*trpBA*). **(c)** JGAB56 was grown alongside WT *A. bescii* over the course of 72 hours at 70 ℃. These two strains were grown three types of defined cellobiose (DC516) media containing 1 mM, 0.1 mM, or no L-tryptophan. Error bars represent one standard deviation from biological triplicates.

The *trpBA* genes encode tryptophan synthase, a heterotetramer consisting of two ⍺-subunits (TrpA) and two β-subunits (TrpB) that catalyze the final step of tryptophan biosynthesis (40, 41). The *trpBA* genes were chosen for deletion (**Fig. 4a**) to minimize disruption to the upstream *trp* biosynthetic genes. Because the open reading frames (ORFs) of *trpF* and *trpB* overlap, the repair template was designed to preserve the portion of the *trpF* ORF overlapping with *trpB*, specifically including the *trpF* sequence through its stop codon in the repair template. Transformation of wild-type *A. bescii* with the Δ*trpBA* CRISPR plasmid (pJLG250), as well as subsequent plating and plasmid curing steps, were carried out as described above for *pyrE* and *ldh*. All colonies picked after the initial plating of transformants at 65℃ yielded PCR amplicons consistent with the expected deletion of *trpBA* (**Fig. S1d & e**). The final sequence-confirmed Δ*trpBA* strain was designated JGAB56 (**Fig. 4b** & **Table 1**). To confirm tryptophan auxotrophy, we compared growth of JGAB56 and wild-type *A. bescii* in defined cellobiose medium (DC516) containing 0, 0.1 mM, or 1 mM tryptophan. Both strains grew similarly with tryptophan supplementation, but only wild-type cells grew without tryptophan supplementation after 72 hours at 70 ℃ (**Fig. 4c**).

Finally, to establish tryptophan-based selection, we constructed replicating vector pJLG253 in which the kanamycin resistance marker (*htk*) was replaced with the *trpBA* genes expressed from the S30 ribosomal protein promoter (P_S30_) (**Fig. 5a**). This vector (pJLG253) also carries a gene encoding the hyperthermophilic β-galactosidase (*Cm*βgal) from the archaeon *Caldivirga maquilingensis*, which we previously developed as a reporter in *A. bescii* (44). This reporter provided a convenient means to demonstrate expression of additional heterologous genes from a vector maintained through *trpBA* complementation. We transformed pJLG253 into the tryptophan-auxotrophic strain JGAB56 (Δ*trpBA*) by electroporation, followed by immediate transfer, without recovery, into selective defined medium lacking tryptophan. Transformant cultures that grew in the tryptophan-free selective medium were plated, and colonies were screened for the presence of the plasmid by PCR (**Fig. S1f**). Sequencing of these PCR products confirmed the presence of the correct P_slp_-*Cm*βgal cassette sequence, and the resulting strain was designated JGAB57 (**Table 1**). To determine whether tryptophan-based selection could support expression of an additional heterologous gene, we measured the activity of the *Cm*βgal reporter in JGAB57 and wild-type *A. bescii* grown in DC516 medium for 30 hours at 70 ℃. JGAB57 exhibited strong reporter activity, whereas no activity was detected in wild-type cells (**Fig. 5b**). These results demonstrate that *trpBA* complementation can be used to select replicating vectors carrying additional heterologous genes in *A. bescii*.

**Figure 5.**
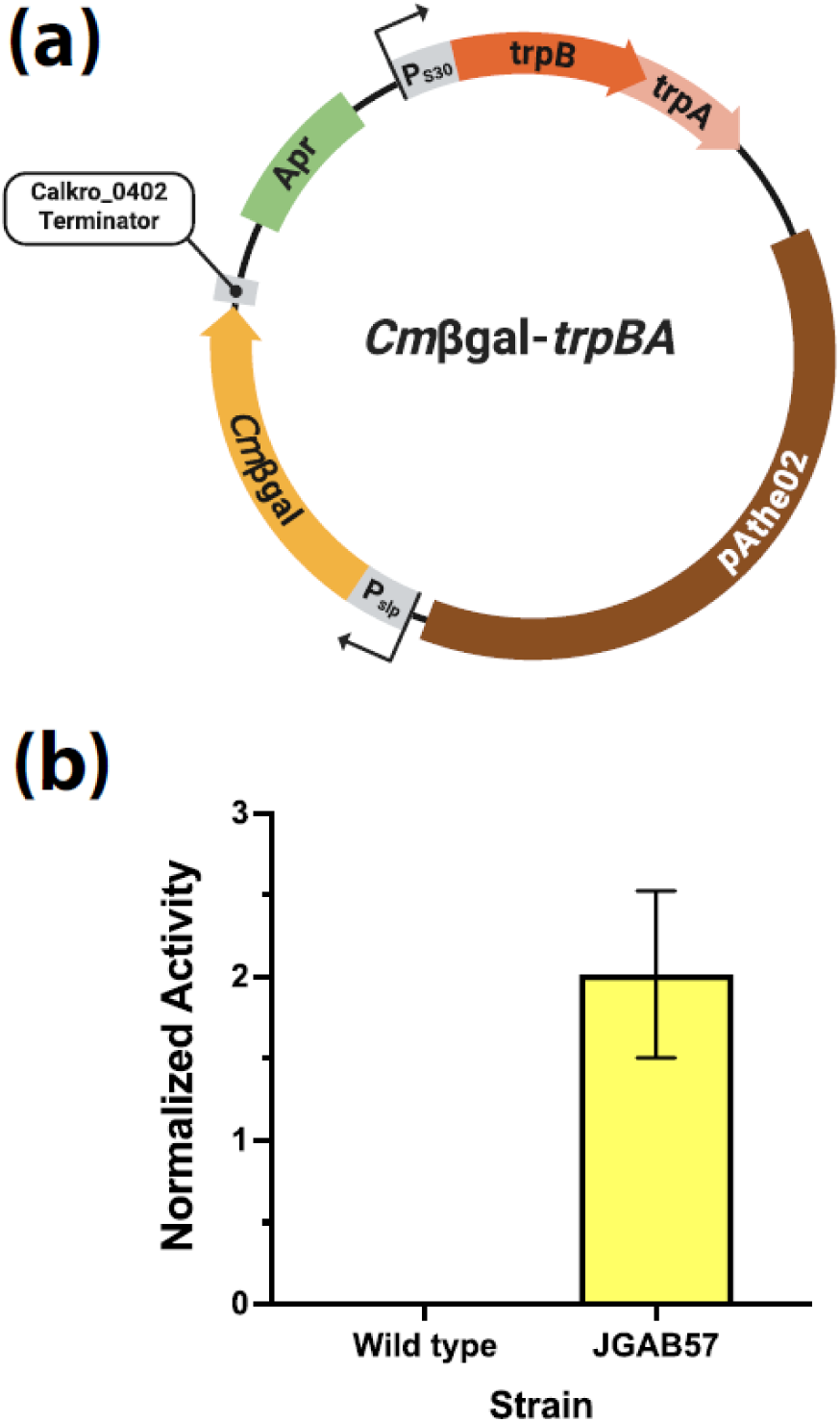
Using *trpBA* complementation for selection in defined media in *A. bescii* strain JGAB56**. (a)** Diagram of replicating vector, pJLG253 (*Cm*βgal-*trpBA*), designed to complement the Δ*trpBA* in JGAB56. The pJLG253 plasmid contains a t*rpBA* complementation cassette for selection of tryptophan prototrophy along with a *Cm*βgal reporter expression cassette. The other elements of this vector’s backbone that enable replication in *E. coli* and *A. bescii* are the same as described above for the CRISPR knock out plasmids. **(b)** Normalized β-galactosidase activity of the wild type *A. bescii* and JGAB57 (*ΔtrpBA* pJLG253) strain after 30 hours of growth at 70 ℃ on DC516 media. Error bars represent one standard deviation among biological triplicates. Panel **a** was created in BioRender. https://BioRender.com/tbqgaos.

Here, we establish the first CRISPR-Cas9 genome editing system in *A. bescii*, enabling targeted and sequential gene deletions through plasmid transformation, plating at 65 °C, and plasmid curing. We further demonstrate the utility of this system by generating a tryptophan-auxotrophic strain and establishing a new plasmid selection strategy based on *trpBA* complementation. Together, these advances expand the genetic toolbox available for *A. bescii* by introducing new approaches to both genome editing and plasmid selection. GeoCas9-mediated editing offers a streamlined alternative to conventional multistep homologous recombination workflows requiring *pyrE/pyrF*-based selection and 5-FOA counterselection. These capabilities have the potential to accelerate the development of engineered *A. bescii* strains for lignocellulose conversion and industrial bioprocessing.

## Materials and Methods

### Cloning and vector construction

Oligonucleotide primers and synthesized DNA used for plasmid construction in this work can be found in **Supplementary Tables S2 & S3** respectively. Full DNA sequences of the five plasmids constructed and used in this study can be found in **Supplementary Table S4**. Maps of the GeoCas9 containing vectors constructed in this study can be found in **Figure 1**, and map of the reporter-complementation vector (pJLG253) can be found in **Figure 5a**. Sequences of gRNAs, repair templates, and the codon optimized GeoCas9 gene can be found in **Supplementary Table S1**. Plasmids were cloned in chemically competent *Escherichia coli* NEB Stable or 10-beta (New England Biolabs). *E. coli* cultures were maintained as previously described with the exception of strains transformed with plasmids containing gRNAs (pJLG240-41 & 50), which were maintained at 28 ℃ for enhanced stability (44). All Gibson assembly steps described below were performed with the NEBuilder HiFi DNA assembly kit (New England Biosciences). Cloned vectors were sequence-confirmed via whole plasmid sequencing (Azenta Genewiz, Plasmid-EZ). Sequence confirmed plasmid DNA was then purified from *E. coli* and prepared for transformation into *A. bescii* as previously described (12, 14, 44).

pJLG236 is a replicating vector that expresses the codon optimized GeoCas9 gene with the *Athe_1331* promoter (P_1331_) but contains no guide RNA (gRNA) or repair template (**Fig. 1**). The backbone of pJLG236 was constructed via PCR linearization (**Table S2**; Primers JLG258 & 286) of a previously constructed vector (pJLG208) that, with P_1331_, expressed the hyperthermophilic β-galactosidase (*Cm*βgal) from the archaeon *Caldivirga maquilingensis* which we developed in a prior work as a reporter in *A. bescii* (37, 44). This backbone consists of a native *A. bescii* replicating plasmid (*pAthe02*), the highly thermostable kanamycin resistance (*htk*) gene expressed by promoter P_S30_ associated with the S30 ribosomal protein (*Athe_2105*), as well as elements for cloning in *E. coli* (e.g. an apramycin marker) (12, 44). The codon optimized gene for GeoCas9 (**Table S1**) obtained as a gift from the lab of Dr. Robert Kelly (North Carolina State University), was amplified (**Table S2**; Primers JLG284-85) with overlaps for subsequent gibson assembly into the pJLG208 backbone. pJLG240, pJLG241, and pJLG250, express gRNAs with a version of the s-layer protein (*Athe_2303*) promoter (P_slp_) truncated at the 3’end to the putative transcriptional start site at the -74 bp position from the native start codon (37); these gRNAs target the genes for lactate dehydrogenase (*ldh*; *Athe_1918*), *pyrE* (*Athe_1382*), and tryptophan synthase (*trpBA*; *Athe_1690-91*), respectively (**Table S1**). Adjacent to the gRNA sequence, these vectors also contain repair templates for homology directed repair consisting of 800 bp flanking regions immediately 5’ and 3’ of the target gene, as well as a unique *PTN* (tagccaacgaattaa) scar site between the two flanking regions (**Table S1**). To assemble these plasmids, pJLG236 was linearized (**Table S2**; Primers JLG292 & 293) at an insertion site between *htk* and the *pAthe02* backbone (**Fig. 1**). For pJLG250, synthesized DNA encompassing both the gRNA and repair template was purchased from Twist Biosciences (**Table S3**). For pJLG240 & pJLG241 only the gRNA and a portion of the repair template could be synthesized by Twist biosciences (**Table S3**), while the remaining part of these repair templates was instead amplified from wild type *A. bescii* gDNA (**Table S2;** Primers JLG295-98). These fragments were synthesized or amplified with appropriate overlaps for subsequent Gibson assembly into the linearized pJLG236 backbone.

pJLG253 is reporter-complementation vector that expresses *trpBA* with P_S30_ for maintenance in Δ*trpBA* strains of *A. bescii* via tryptophan prototrophy on defined media and also contains the *Cm*βgal reporter gene expressed by P_slp_ (**Fig. 5a**). pJLG253 was constructed from another previously constructed reporter plasmid (pJLG093) that expressed *Cm*βgal with P_slp_ but contained *htk* as the selection marker (44). To construct pJLG253, pJLG093 was first linearized (**Table S2;** Primers JLG323 & 24) as to remove the gene for *htk* while leaving P_S30_ intact. The *trpBA* gene cassette was then amplified with appropriate overlaps from WT *A. bescii* gDNA (**Table S2;** Primers JLG325 & 26), and Gibson assembled into the pJLG093 backbone.

### A. bescii growth and maintenance

*A. bescii* strains were cultured in sealed serum bottles or “Balch-type” anaerobic tubes, both of which were sealed with 20 mm butyl rubber stoppers. Unless described otherwise, cells were grown at 70-72 ℃ without shaking in C516 medium following the recipe described by *Lipscomb et al.* with the carbon source at 5 g/L indicated by a second letter of the medium identifier (e.g. CM516 medium contains maltose) and the following additional ingredients: 0.5 g/L yeast extract, 1x modified DSM516 salt solution, 1x trace element solution, 1x vitamin solution, 0.25 µg/L resazurin, 0.16 µM sodium tungstate, 1 g/L cysteine hydrochloride, 1 g/L sodium bicarbonate, and 1 mM potassium phosphate buffer pH 7.2 (12). Stock solutions of the 200x vitamin and 1000x trace element solutions were prepared as described previously (1, 12). The modified 50x stock salt solution contained 16.5 g/L MgCl_2_•6H_2_O, 16.5 g/L KCl, 16.5 g/L NH_4_Cl, and 7 g/l CaCl_2_•2H_2_O. The defined (D516) version of this media excludes yeast extract. As needed, C516 medium was further supplemented with 50 µg/ml kanamycin (IBI Scientific), 20 µM uracil (TCI Chemicals), and 1 mM or 0.1 mM of L-tryptophan (TCI Chemicals). Sealed bottles or tubes containing filter sterilized media were made anaerobic through gas sparging, with the headspace being replaced with 80% (v/v) N_2_ and 20% (v/v) CO_2_ gas. Cell density of *A. bescii* cultures was measured as the optical density at 680 nm (OD680) using a cuvette in a Nanodrop One C spectrophotometer (Thermo Scientific) blanked with 1x modified DSM 516 salt solution.

### Genetic manipulations of *A. bescii*

A table of strains constructed in this study can be found in **Table 1**. The parent strain for JGAB51, JGAB53, JGAB54, and JGAB56 was wild type *A. bescii* DSM 6725 obtained from the lab of Dr. Robert Kelly (North Carolina State University). JGAB55 was constructed from JGAB54, while JGAB57 was constructed from JGAB56.

For construction of JGAB51, and JGAB53-55, *A. bescii* were transformed as previously described with competent cells grown on maltose media (CM516) supplemented with amino acids (12, 14, 44). Briefly, competent cells were first washed in 10% (w/v) sucrose and then transformed via electroporation in 1mm cuvettes using a Bio-Rad gene pulser (1800-2200 V, 400-600 Ω, and 25 µF) with plasmids previously methylated *in vitro* using the M.CbeI methyltransferase. After 1-2 hours of recovery at 70-72 ℃, transformants were transferred to CM516 media containing kanamycin for selection. Uracil at 20 µM was also supplemented in media throughout recovery and selection for construction of JGAB53-55. JGAB56 was transformed similarly except recovery, and selective media were supplemented with 1 mM tryptophan. To construct JGAB57, competent JGAB56 cells were grown in CM516 media supplemented with amino acids and additional tryptophan such that the final concentration of tryptophan was 1 mM for consistency. Washing and electroporation steps were performed as described above, after which cells were immediately transferred without a recovery step into 50 ml of pre-heated defined maltose media (DM516) for selection via tryptophan prototrophy. Cultures from the above transformations that grew in selective media were subsequently plated as described below in an anaerobic chamber under a >98% (v/v) N_2_ and 1-2% (v/v) H_2_ atmosphere.

For JGAB51, transformants were plated and grown in cellobiose (CC516) media at 70 ℃, after which single colonies were picked into the same media. The procedure was similar for JGAB57 except cells were grown in defined cellobiose (DC516) media without kanamycin. Picked colonies were then screened via colony PCR (**Table S2;** Primers JLG181-82 & 323). The presence of the correct GeoCas9 sequence in JGAB51 and P_slp_-*Cm*βgal reporter sequence in JGAB57 were confirmed by long-read sequencing of the colony PCR products (Azenta Genewiz, PCR-EZ).

For construction on JGAB53-56, which were transformed with gRNA containing CRISPR vectors, transformants were also initially plated and grown in solid CC516 media with kanamycin but at lower temperature of 65 ℃ to facilitate GeoCas9 activity. CC516 media was additionally supplemented with uracil or tryptophan as appropriate throughout the process of generating these strains. During construction of JGAB53 and JGAB54, which represented the initial implementation of CRISPR in *A. bescii*, transformants were also plated alongside cultures of JGAB51 as a negative control. Single colonies were picked into the same medium as plating and screened for the desired genetic knockouts via colony PCR (**Table S2;** Primers JLG288-91 & 319-20) outside the flanking regions in the repair templates. The presence of the correct genomic deletion and incorporation of the *PTN* scar site was confirmed by long-read sequencing of the colony PCR products (Azenta Genewiz, PCR-EZ). PCR was also routinely performed on the initial transformant liquid cultures prior to plating to monitor their genotype status. Next, to purify the edited strains and cure them of the CRISPR editing vectors, sequence confirmed colonies were re-plated in non-selective (no kanamycin) CC516 media at 70 ℃, after which individual colonies were then picked into the same non-selective medium. This process was repeated one more time, after which the final picked colonies were re-sequence confirmed for the correct genotype. Finally, to test if they were cured of their CRISPR plasmids, final purified strains were grown for at least 40 hours at 70 ℃ on CC516 media with or without kanamycin alongside a kanamycin resistant strain of *A. bescii* as a positive control. No growth was detected on kanamycin media for any of the plate purified strains, indicating they had been cured of their CRISPR plasmids.

### Characterization of generated *A. bescii* strains

Growth experiments testing for uracil or tryptophan auxotrophy in *A. bescii* were conducted over the course of 72 hours, with OD680 cell density measurements taken once every 24 hours. Cells were grown in bottles at 70 ℃ in defined cellobiose media (DC516). To begin growth curves, inoculum cultures were first washed in sugar-free defined media (D516) as described by *Tjo et al* (45). These washed cells were then used to inoculate pre-heated media to a target OD680 of ∼0.005, with cells in all conditions being grown in biological triplicate. To test if the JGAB54 (*ΔpyrE*) strain was an uracil auxotroph, it was grown alongside wild type *A. bescii* as described above in DC516 media with or without the addition of 20 µM uracil. Inocula for both strains were grown previously in DC516 media supplemented with 20 µM uracil. The necessity of tryptophan for the growth of JGAB56 (*ΔtrpBA)* was similarly tested against wild type *A. bescii* by growing the two strains on DC516 media without tryptophan, and the same media supplemented with 1 mM or 0.1 mM tryptophan. Inocula for this latter experiment were grown previously in DC516 media supplemented with 1 mM tryptophan.

To determine if lactate production was eliminated in JGAB53 (*Δldh*), this strain was grown alongside wild type *A. bescii* in tubes of DC516 media (no uracil). To begin the experiment, pre-heated media was inoculated to a target OD680 of ∼0.005 with cells previously grown in the same DC516 media. Cultures were then grown in biological triplicate for 30 hours at 70 ℃, after which final OD680 measurements were taken, and additional samples were removed for analysis. Lactate and acetate produced from fermentation were quantified via comparison to standards on an HPLC. Samples were prepared for HPLC via centrifugation at 21000 x g for 5 minutes followed by filtration of the supernatant through a 0.22 µm filter. Samples were analyzed alongside a filtered blank of DC516 media using a 300 x 7.7 mm Agilent Hi-Plex H column heated to 50 ℃, with a mobile phase of 5 mM H_2_SO_4_, a flow rate of 0.6 mL/min, and an injection volume of 10 µL. Data was collected with an Agilent 1260 Infinity II Refractive Index detector also heated to 50 ℃.

To test if JGAB57 (*ΔtrpBA* pJLG253[*trpBA, Cmβgal*]) expressed the *Cm*βgal reporter, JGAB57 and wild type *A. bescii* were inoculated and grown in DC516 media as described above for the experiment testing JGAB53. At the end of growth, OD680 measurements were taken, and 5 ml samples of culture were removed and harvested for subsequent reporter assays as previously described (44). Reporter activity in *A. bescii* was assayed as described previously for *Cm*βgal (44). Briefly, β-galactosidase activity was detected using the colorimetric substrate para-nitrophenyl-β-D-galactopyranoside (pNPβGal) obtained from TCI chemicals, with the substrate solution containing 5 mM pNPβGal dissolved in pH 5.5 100 mM sodium acetate pH 5.5 buffer. Harvested *A. bescii* cells were prepared in 100 mM pH 5.5 sodium acetate buffer, heat-treated for 10 minutes at 90 ℃, after which they were added to the substrate solution and incubated for another 10 minutes at 90 ℃. Reactions were then quenched with addition of 1M sodium carbonate, and the absorbance at 405 nm (A405) of each reaction was measured in a flat-bottomed clear 96 well plate using an Infinite® M Plex (Tecan) microplate reader. Normalized activity was also calculated as described previously with normalization by the OD680 of the prepared *A. bescii* cells input into each reaction (44). All reaction conditions, including for each biological replicate, were performed in technical triplicate.

## Data Availability Statement

The data associated with this study are available in the article and its supporting information.

## Acknowledgements

This work is supported by the A1531 Biorefining and Biomanufacturing program, project award no. 2026-67022-45882, from the U.S. Department of Agriculture’s National Institute of Food and Agriculture, as well as startup funds from the Department of Chemical and Biological Engineering at Princeton University. We would also like to acknowledge Hansen Tjo for his helpful advice regarding the growth and maintenance of *A. bescii*.

## Conflict of Interest

J.L.G. and J.M.C. are inventors on a provisional patent application related to genetic engineering of *A. bescii*.

